# Vitamin A is necessary for acquisition, but not for expression or progression, of CNS autoimmunity

**DOI:** 10.1101/2025.05.18.654726

**Authors:** Reiko Horai, Ru Zhou, So Jin Bing, Kaska Wloka, Yingyos Jittyasothorn, Todd E Duncan, Mary J Mattapallil, Phyllis B Silver, Chi-Chao Chan, Rachel R Caspi

## Abstract

Vitamin A (VitA) and its derivative retinoic acid (RA) are essential for immunological responses. In VitA deficient (VAD) mice, acquisition of effector responses is impeded, but little is known about maintenance and expression of previously acquired effector function under the VAD conditions. We examined the impact of VAD on progression of autoimmune diseases using two models of uveitis, experimental autoimmune uveitis (EAU) induced by active immunization and spontaneous uveitis in retina-specific T cell receptor transgenic (R161H) mice, and in the model of experimental autoimmune encephalomyelitis (EAE). VAD was induced by dietary lack of VitA from before birth, or by daily injections of a pan-RA receptor inhibitor BMS493 in adult mice fed with the standard diet. VAD mice were essentially resistant to induction of EAU or EAE and displayed impaired effector T cell responses. Defective priming/acquisition of effector function by VAD T cells was also evident. By contrast, spontaneously uveitic R161H mice fed with VAD diet, in which priming of pathogenic T cells occurs before onset of full VAD, only moderately attenuated uveitis compared to VitA sufficient R161H mice. To reconcile somewhat different results between induced model and spontaneous model of uveitis, we examined EAU in partial VAD mice or adoptive transfer into VAD hosts. The results supported that effector T cells primed in VitA-sufficient environment were able to function in VAD environment and induced EAU. We conclude that although priming of naïve T cells in the VAD environment is defective, effector function acquired under VitA sufficient conditions is maintained and can be expressed under VAD conditions. Because dietary lack of VitA is rarely profound and may be seasonal, our findings may shed light on immunity and autoimmunity in geographical regions where dietary VitA is limiting.

## INTRODUCTION

Vitamin A (VitA) and its metabolites are essential for major biological functions, including development, vision and immunological responses. VitA maintains the integrity of both innate and adaptive immune system, mainly through its active metabolite *all-trans* retinoic acid (RA), which is critical for the homeostasis of T helper cells, particularly in mucosal immunity. It is important for imprinting CD4^+^ and CD8^+^ T lymphocytes as well as B cells to traffic to the gastrointestinal mucosa [1-4]. RA produced by intestinal DC synergizes with IL-6 or IL-5 to induce IgA production [4, 5]. Arguably the best-known function of RA in immunity is in generation of induced T regulatory cells (Tregs) in the presence of TGF-β, while at the same time inhibiting Th17 cell differentiation [6-9]. This has implications for control of autoimmunity, as demonstrated by reports that retinoid treatment ameliorates a number of experimental autoimmune diseases in rodents as well as colitis in the human [10-15]. Due to its role in Treg induction and T cell trafficking and expansion in the gut, RA is also involved in induction the process of oral tolerance [16, 17]. In the eye, RA is naturally present in ocular fluids as one of the integral components in the normal visual cycle [18] and contributes to ocular immune privilege by inducing Tregs *in vivo* locally within the eye [19, 20]. Although RA has been studied mainly as an anti-inflammatory molecule balancing the Treg and Th17/Th1 immunity, other studies uncovered that it is also required to for effector T cell activation and acquisition of adaptive CD4 immunity to infection and mucosal vaccination [21]. Additionally, in the presence of IL-15, RA acts as co-adjuvant to induce Th1 immunity to dietary antigens, resulting in the breakdown of tolerance to gliadin in the intestine in a mouse model of coeliac disease [22].

Complete and protracted VitA deficiency (VAD) is probably rare even in the developing world. Rather, it could fluctuate depending on seasonal changes in nutrition, and partial or insufficient VAD (VitA insufficiency; VAI) may be more reflective in the real world. In this sense, the studies of Hall et al. and DePaolo et al. have concentrated on the role of VitA in T cell activation and acquisition of adaptive immunity [21, 22], but little is known about maintenance and expression of previously acquired effector function under conditions of VAD or VAI. Also still unaddressed are effects of VAD on autoimmune disease, where effector cells can persist for a long time and their target antigen is always present, as these studies concentrated on host defense and dietary antigen. Our current study addresses these gaps in knowledge in animal models of central nervous system (CNS) autoimmunity: (1) Experimental autoimmune uveitis (EAU) is an induced autoimmune uveitis model, driven by a response to retinal autoantigen, interphotoreceptor retinoid binding protein (IRBP), following active immunization. (2) R161H mice express a transgenic T cell receptor (TCR) specific for IRBP and serve as a spontaneous model of autoimmune uveitis affecting the neuroretina [23]. Both EAU and R161H represent human autoimmune uveitis which is estimated to underlie 10-15% of the cases of severe visual handicap [24, 25]. (3) Experimental autoimmune encephalomyelitis (EAE) is a well-studied induced autoimmune disease model for human multiple sclerosis, affecting the CNS.

Our results demonstrate that EAU induced by immunization under conditions of VAD is strongly attenuated due to deficits in T cell differentiation and effector function. Similar results were supported by EAE experiments. In contrast, autoimmunity to the neuroretina in the R161H spontaneous model, in which autoimmune effector priming starts before the onset of complete VAD, disease is maintained and only marginally attenuated under the VAD condition. Notably, effector function and development of pathology are maintained despite the VAD state in the host when EAU was induced and autoreactive T cells were primed prior to the full VAD state, or when the retina-specific T cells were activated in vitro and adoptively transferred into the VAD recipients. These results lead us to conclude that although acquisition of effector function by naïve T cells under VitA deficiency is defective, effector function that was acquired previously is fully maintained in the VAD host and can drive autoimmune disease. These findings may have clinical implications in geographical regions with insufficient VitA intake.

## MATERIALS AND METHODS

### Mice and VAD induction

B10.RIII mice were obtained from Jackson Laboratory (Stock number 000457). An IRBP-specific TCR transgenic mouse line (R161H) on the B10.RIII background was developed in house [23]. Foxp3^GFP^ reporter mice (a kind gift from Dr. A. Rudensky, Sloan-Kettering Cancer Ctr, New York, NY) [26] were backcrossed onto the B10.RIII background. The two strains were crossed to produce R161H-Foxp3^GFP^ reporter mice and were used for monitoring the spontaneous uveitis and for cell donors. Diet-induced VAD mice were produced as described [1]. Briefly, pregnant dams were placed on a diet lacking Vit A (cat# TD.09838, Envigo, Madison, WI) as soon as pregnancy could be detected at day 14 of gestation. Control mice were fed the same diet supplemented with 20,000 IU vitamin A /kg (cat# TD.09839). Pups were maintained on the same diet after weaning and were used after 9 weeks of age. In some experiments, a pan-RA receptor inhibitor BMS493 was used to mimic VAD conditions in adult mice fed with the regular facility diet (contains 24,200 IU VitA /kg, NIH-31 7017). Mice were injected i.p. with daily doses of either vehicle (DMSO) control or BMS493 (220 µg/day)[27]. Other mice (e.g., cell donors) were used at 6-12 weeks of age. All mice were housed under specific pathogen-free conditions. Treatment of animals was in compliance with Institutional Guidelines, and animal study protocols, NEI-583 (EAE) and NEI-688 (EAU), were approved by the National Eye Institute Institutional Animal Care and Use Committee.

### Measurement of retinol levels

The level of VitA was determined by using an adapted high performance liquid chromatography (HPLC) method [28]. The retinol in serum or eye homogenate was extracted into hexane, dried under gentle nitrogen stream, and re-dissolved in methanol, and run through a Poroshell 120 EC-C18 reversed-phase column (4.6 x 50 mm, 2.7 μm; Agilent). Retinyl acetate was added as an internal standard to confirm extraction efficiency. Data were analyzed using Empower 3 software (Waters Corp., Milford, MA). All procedures involving retinoids were conducted under gold light.

### Visual function assessment by electroretinogram (ERG)

Visual function of the eyes was evaluated by dark- and light-adapted ERG using Espion E2 System (Diagnosys LLC). Mice were dark adapted overnight, and all procedures were performed under dim red light. Mice were anesthetized by intraperitoneal injection of ketamine-xylazine (77 mg/kg and 4.6 mg/kg, respectively), and their pupils were dilated using Tropicamide and phenylephrine (0.5%). Electrodes were placed on center of the cornea, a reference electrode was placed inside the mouth, and a ground electrode was inserted under the back skin. The a-wave amplitude was measured from the baseline to the trough of the a-wave and b-wave amplitude was measured from trough of the a-wave to the peak of the b-wave.

### CD4^+^ T cell isolation and culture

CD4^+^ T cells were purified from pooled splenocytes and/or lymph nodes, either by sorting for CD4^+^GFP^-^ T cells to ∼99% purity using FACS Aria (Becton Dickinson, San Jose, CA), or by immunomagnetic isolation (Miltenyi Biotec, Auburn, CA) for CD4^+^CD25^-^ T cells (residual Foxp3^+^ T cells ∼1.5%). Cells in complete HL-1 medium (Cambrex, East Rutherford, NJ) were stimulated with plate-bound anti-CD3 Ab (5μg/ml) plus soluble anti-CD28 Ab (2 μg/ml) for 36 h (for RNA) or 3 days (for ELISA). Where indicated, cells were polarized under Th1-polarizing conditions (10 ng/ml mIL-12, 10 µg/ml anti-IL-4), Th17-polarizing conditions (0.5 ng/ml hTGF-β1, 10 ng/ml mIL-6, 10 µg/ml anti-IFN-γ, 10 µg/ml anti-IL-4), or Treg-inducing conditions (5 ng/ml hTGF-β, 10 ng/ml hIL-2). Recombinant cytokines were from R&D Systems (Minneapolis, MN), anti-IFN-γ (clone R4-6A2) was from BioXcell (West Lebanon, NH), and anti-IL-4 (11B11) was from National Cancer Institute-Frederick Biological Resources Branch Preclinical Repository (Frederick, MD). Cytokine concentrations in culture supernatants were determined by ELISA (R&D).

### Treg suppression assay

CD4^+^GFP^+^ regulatory T cells (Tregs) from control or VAD Foxp3^GFP^ reporter mice were sorted to purity (see above) and co-cultured at various ratios with sorted CD4^+^GFP^-^ or CD4^+^GFP^-^ CD44^high^ (memory/activated) responder T cells (5×10^4^cells/well) from Foxp3-GFP reporter mice or R161H Foxp3-GFP reporter mice in the presence of soluble anti-CD3 Ab (0.5μg/ml) or human IRBP_161-180_ peptide (1 μg/ml), respectively, and irradiated T cell-depleted spleen cells (1×10^5^cells/well) for 72 h. [^3^H]-thymidine (1μCi/well) was added for the last 8-12 h of culture. The incorporated radioactivity was measured by a MicroBeta TriLux scintillation counter (PerkinElmer Life Science, Boston, MA, USA). To examine the IL-2 and IL-10 gene expressions, responder cells were harvested 36 h after co-culture as described above, and sorted for CD4^+^GFP^-^ responders, followed by real-time PCR analysis (described below).

### Flow cytometric analysis

For cell surface staining, cells were stained with the indicated antibodies for about 30 min, followed by washes with PBS containing 2% FBS. Then cells were resuspended, stained with cell viability dye, and analyzed by flow cytometry. For intracellular cytokine staining, cells were stimulated with PMA (10 ng/ml) and Ionomycin (500 ng/ml) in the presence of Brefeldin A (GolgiPlug, BD Biosciences, San Diego, CA) as described elsewhere. After 4 h, cells were fixed with 4% paraformaldehyde, permeabilized with PBS containing 0.1% BSA and BD Wash/Perm buffer (BD) and stained with the indicated antibodies. Antibodies used for flow cytometry analysis and cell sorting were purchased from BD Biosciences, Invitrogen (San Diego, CA) or BioLegend (San Diego, CA). Up to 100,000 events were acquired on BD FACSCalibur, BD FACS Aria II, MACSQuant Analyzer 10 (Miltenyi, Bergisch Gladbach, Germany), or CytoFLEX (Beckman Coulter, Indianapolis, IN), depending on the experiment. Data were analyzed using FlowJo software (TreeStar Inc, Ashland, OR).

### RNA isolation and Real-time PCR

Total RNA from cells was extracted with RNeasy Mini Kit (QIAGEN, Valencia, CA) and cDNA was synthesized using SuperScript III First-Strand Synthesis System (Invitrogen). Quantitative real-time PCR (qRT-PCR) was performed with TaqMan 7500 sequence detection system (Applied Biosystems, Foster City, CA) using GAPDH for endogenous control and primer/probe sets from Applied Biosystems. CD4^+^ T cells from pooled spleens and lymph nodes were prepared from naïve B10.RIII Foxp3^GFP^ reporter mice, and their baseline gene expression levels were used as naïve control. Data were normalized to respective GAPDH and expressed relative to naive control (set as 1).

### Induction and scoring of EAU by immunization or adoptive transfer

To induce EAU by active immunization, mice were subcutaneously injected with human IRBP161-180 peptide (5-6 µg/mouse) emulsified in complete Freund’s adjuvant (CFA, Difco, Detroit, MI) that had been supplemented with *Mycobacterium tuberculosis* strain H37RA to 2.5 mg/ml, as described [29]. To induce EAU by adoptive transfer, peripheral lymph node cells collected from R161H mice were stimulated with 2 μg/ml hIRBP161-180 peptide for 3 days under Th1-polarizing conditions (10 ng/ml IL-12, 10 µg/ml anti-IL-4; 10 ng/ml IL-2 was added on day 2 of culture) or Th17-polarizing conditions (2 ng/ml TGF-β, 10 ng/ml IL-6, 10 µg/ml anti-IFN-γ, 10 µg/ml anti-IL-4; 10 ng/ml IL-23 was added on day 2 of culture). Live cells were isolated by Lympholyte^®^-M (CEDARLANE, Burlington, NC) and injected intraperitoneally into control or VAD recipients.

Disease scores were read in a masked fashion by fundus examination and confirmed by histopathology on eyes collected at the termination of the experiment (20-21 days after immunization or 11-14 days after adoptive transfer). Clinical scores by fundoscopy were assigned on a scale from 0 (no inflammation) to 4 (complete destruction of the retina) in half-point increments as described previously [30]. Histopathology of the eyes was performed on methacrylate-embedded sections stained with hematoxylin and eosin, and the severity of EAU was evaluated on a scale of 0-4 based on the number, type, and size of lesions as described previously [30, 31].

### Induction and scoring of EAE

B10.RIII WT mice were immunized with 300 μg of whole MBP in CFA that had been supplemented with Mtb strain H37RA to 4 mg/ml. As an additional adjuvant, 0.3 µg pertussis toxin (Sigma-Aldrich, St. Louis, MO) was i.p. injected on day 0 and day 2 [32]. Clinical assessment of EAE: 0, no clinical signs; 1, flaccid tail; 2, hind limb weakness or abnormal gait; 3, complete hind limb paralysis; 4, complete hind limb paralysis with forelimb weakness or paralysis; and 5, moribund or deceased.

### Depletion of CD25^+^ T cells by PC61 treatment

To deplete CD25^+^ cells (representing mostly regulatory T cells in naïve mice), animals were given 2 i.p. injections, 3 days apart, of 1.0 mg of the anti-CD25 mAb, PC61 [33, 34]. Six days after the second injection, depletion was determined by flow cytometry in peripheral blood mononuclear cells (PBMC) of representative animals and all mice were immunized for EAU on the following day. Depletion of CD25^+^ cells was observed as reduction of CD4^+^CD25^+^GFP(Foxp3)^+^ cells stained with anti-CD25 mAb (Clone 7D4), which binds to a different epitope than PC61 on the IL-2 receptor [35]. Mice were then immunized with a low dose of IRBP161-180 (2.5 µg/mouse) expected to elicit mild to moderate disease scores, so as to demonstrate enhanced disease in depleted *vs.* non-depleted mice.

### Intravitreal injection and analysis of retina-specific donor T cells

CD4^+^GFP^-^ T cells were purified from lymph nodes and spleens of R161H-Foxp3^GFP^ mice expressing CD90.2 on a conventional (non-Rag-deficient) background, with ∼30% TCR transgenic T cells. About 0.5 million cells were injected through the pars plana into the posterior chamber of each eye of CD90.1 congenic recipients. Eyes that sustained unintentional damage to the lens or other structures as a result of the injection were excluded from analysis. Six to seven days after cell injection, disease was scored by fundoscopy. Recipient eyes were harvested and dispersed into single cell suspension as described [36].

### Statistics and experimental reproducibility

Values are presented as mean ± SEM where indicated. Statistical differences were calculated with an unpaired Student’s *t*-test, two-tailed (GraphPad Prism versions 7-10). Statistical significance was set at p ≤ 0.05. Experiments were repeated at least two, and usually more three times, with number of repetitions stated in the figure legends. Groups numbered 3–7 mice, depending on the experiment (usually 4-5).

## RESULTS

### The process of VAD induction does not compromise visual function or T cell development

VAD by dietary induction was verified by measuring serum levels of retinol, an alcohol form of VitA, by HPLC. In the VAD mice at 9 weeks or older, the serum retinol levels were significantly lower than age-matched VitA-sufficient control mice (Fig. 1A). Relative serum retinol levels in mice on the VAD diet were already less than 40% of those on the control diet at 5 weeks of age, and show further gradual reductions until 9 weeks of age to become “VAD” at that point (Fig. 1B).

**Figure 1.**
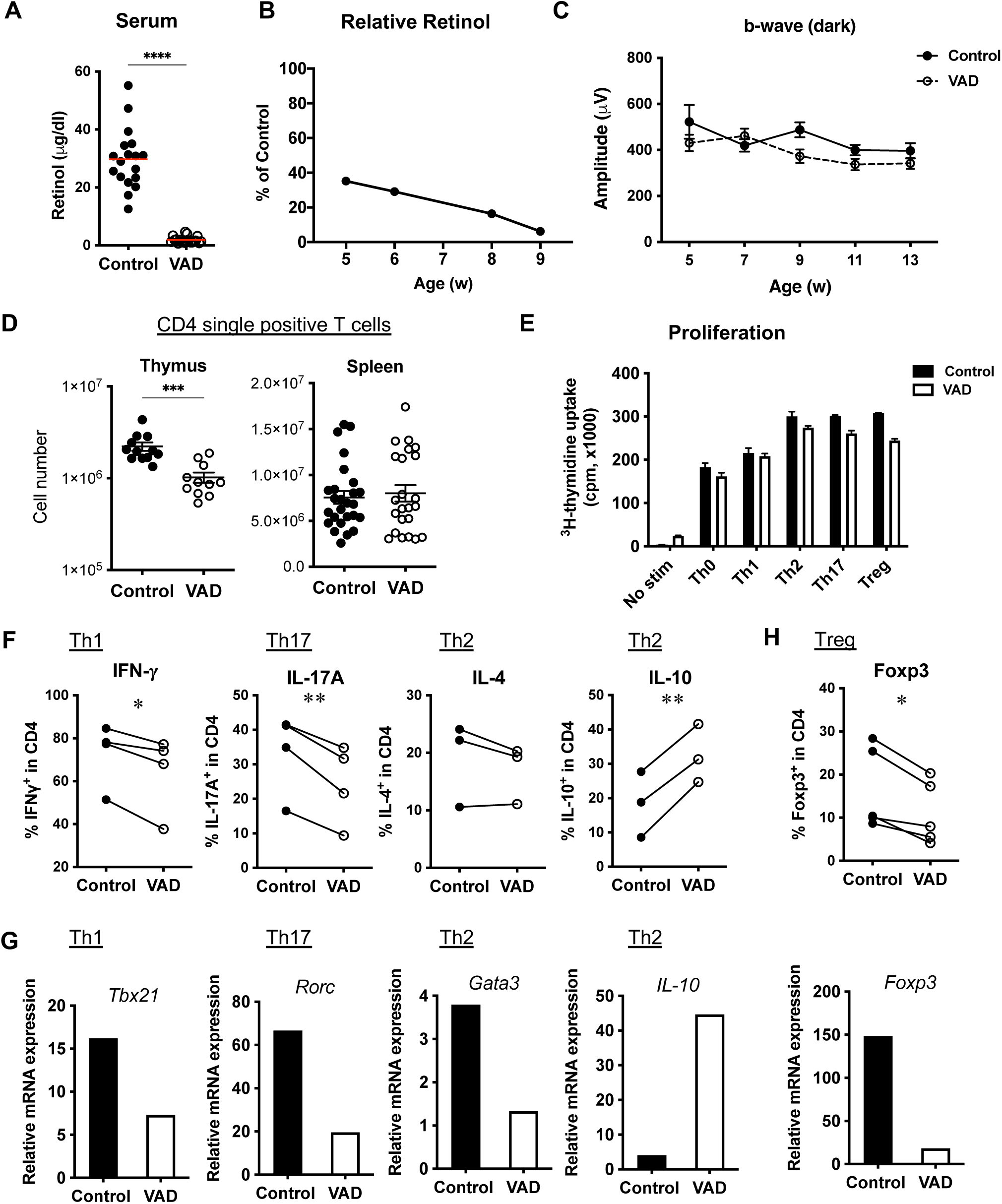
VAD mice develop normal visual function and peripheral T cells, but show impaired Th differentiation in vitro. (**A**) VAD was confirmed by measuring retinol levels in serum by HPLC. \*\*\*\**P*<0.0001. (**B**) Relative serum levels of retinol in mice on the VAD diet to controls show gradual reductions by age. (**C**) Visual function measured by electroretinography (ERG) on naïve B10.RIII WT mice was not impaired. Mice were dark-adapted overnight prior to the recording. Data include younger ‘partial’ VAD stage (5-, 7-week-old) and VAD after 9-week-old. Representative one experiment of three is shown. N=13 for control mice, and N=18 for VAD mice. (**D**) CD4 T cell numbers in the thymus and spleen of VAD mice. Data are compiled from 6 to 8 experiments. \*\*\**P*=0.001 by Mann-Whitney test. (**E-H**) Splenic CD4^+^CD25^-^ T cells isolated from unimmunized VAD or VitA-sufficient mice were activated with plate-bound anti-CD3 and soluble anti-CD28 Abs for 36 h (for mRNA levels measured by RT-PCR), 48 h (for proliferation) or 3 days (for protein detection by flow cytometry) in vitro under Th1, Th17, Th2 or Treg conditions. (**E**) Proliferation measured by [^3^H]-thymidine uptake. One representative data of two experiments. (**F**) Intracellular cytokines detected by flow cytometry under respective conditions. Paired data of control vs VAD from 3-4 experiments are shown with connected lines. \**P*<0.05, \*\**P*<0.01 by paired t-test. (**G**) mRNA expression of Th-relevant transcription factors and IL10 (Th2) under the respective conditions was determined by q-PCR. One representative experiment of two or more is shown. (**H**) Foxp3 expression under Treg conditions was analyzed by flow cytometry and RT-PCR. Data are representative of 5 experiments.

Since VitA is important for maintaining visual function, and its deficiency can cause night blindness and dry eyes in humans [37], we examined visual function of experimental mice by electroretinogram (ERG). Fig. 1C shows that visual function was not impaired in mice on the VAD diet at all the time points examined. This is expected when they were still in the process of becoming VAD (5-7 weeks old), but was surprising that the visual function was unaffected by VAD after 9 weeks of age, when animals were highly VitA-deficient. The results indicate that visual function of our experimental VAD mice is relatively normal.

To investigate whether VAD affects the T cell development *in vivo*, we compared lymphocyte cellularity in thymus and spleen between control and VAD mice. Total numbers of lymphocytes were comparable in thymus and spleen (Suppl. Fig. S1). There was a significant reduction of CD4 single positive thymocytes in VAD mice compared to control mice, but the CD4^+^ T cell cellularity in VAD mice was restored in the periphery and appeared normal in spleen (Fig. 1D). These results suggest that the effects of VAD induction on T cell development are marginal and that these T cells and mice can be useful for *in vitro* and *in vivo* studies on T cell-mediated autoimmunity.

### T cells from VAD mice show impaired effector T cell differentiation *in vitro*

To examine T cell proliferation to antigenic exposure in VAD mice, CD4^+^CD25^-^ T cells from spleens were isolated as a naïve population and stimulated with anti-CD3/CD28 Abs. VAD T cells proliferated similarly to control T cells under the non-polarizing (Th0) condition or Th-polarizing conditions (Fig.1E), indicating that T cell proliferation via TCR activation is not affected by VAD. To examine if T cell differentiation is impaired in VAD mice, CD4^+^CD25^-^ naïve T cells were cultured under Th1, Th17, Th2 or Treg polarizing conditions for 36-72 h and the expression of representative lineage-specific cytokines and transcription factors was assessed by real-time q-PCR or flow cytometry (Fig. 1F-H). Notably, under both Th1- and Th17-polarizing conditions, VAD CD4^+^ T cells showed significantly decreased frequencies of IFN-γ or IL-17A-producing populations, respectively (Fig. 1F). The data was supported by the lower expression of transcriptomes for T-bet (*Tbx21* for Th1) and RORγt (*Rorc* for Th17) (Fig. 1G).

On the other hand, under Th2-polarizing conditions, IL-4-producers were not affected, despite the lower expression of *Gata3* mRNA, while IL-10 expressing cells were rather enhanced in VAD CD4^+^ T cells (Fig. 1F, G). Furthermore, under Treg-inducing conditions, Foxp3 expression was reduced both in the protein and transcriptome levels (Fig. 1H). These results indicate that T cells, upon reaching the VAD status, have defective capacity of differentiation into Th1 and Th17 effectors or into regulatory cells.

### Tregs isolated from VAD mice are predominantly Helios+ and are functionally capable of inhibiting activation of naïve T cells (Figure S1 – development and function of Tregs)

We next examined the Treg compartment of VAD mice. In the thymus, total Foxp3^+^ cell counts and frequencies within CD4^+^ cells were comparable between control and VAD mice (Fig. 2A). Helios transcription factor and neuropilin-1 serve as markers for thymically derived Tregs (nTregs) in unmanipulated mice [38, 39]. Helios expression in nTregs in thymus was unchanged, but neuropilin-1 expression was significantly decreased in VAD mice. In spleen, total Foxp3^+^cell counts and frequencies within CD4^+^ T cells were increased in VAD mice (Fig. 2C). Staining for Helios revealed that Foxp3^+^ T cells of VitA-sufficient mice contained both Helios-positive nTreg and Helios-negative peripherally induced Treg (Figure 2B). In contrast, Foxp3^+^ cells from VAD mice were composed of mostly Helios-positive (nTreg) cells. Together with the impaired conversion of VAD T cells to Tregs *in vitro* (Fig. 1H), these data are compatible with the interpretation that *de novo* induction of Tregs in the periphery is reduced in the VAD environment, resembling defective acquisition of effector function by non-Tregs at that stage. However, the effects on nTregs are minor, because the nTreg development in the thymus has been largely completed before the VAD was achieved *in vivo*.

**Figure 2.**
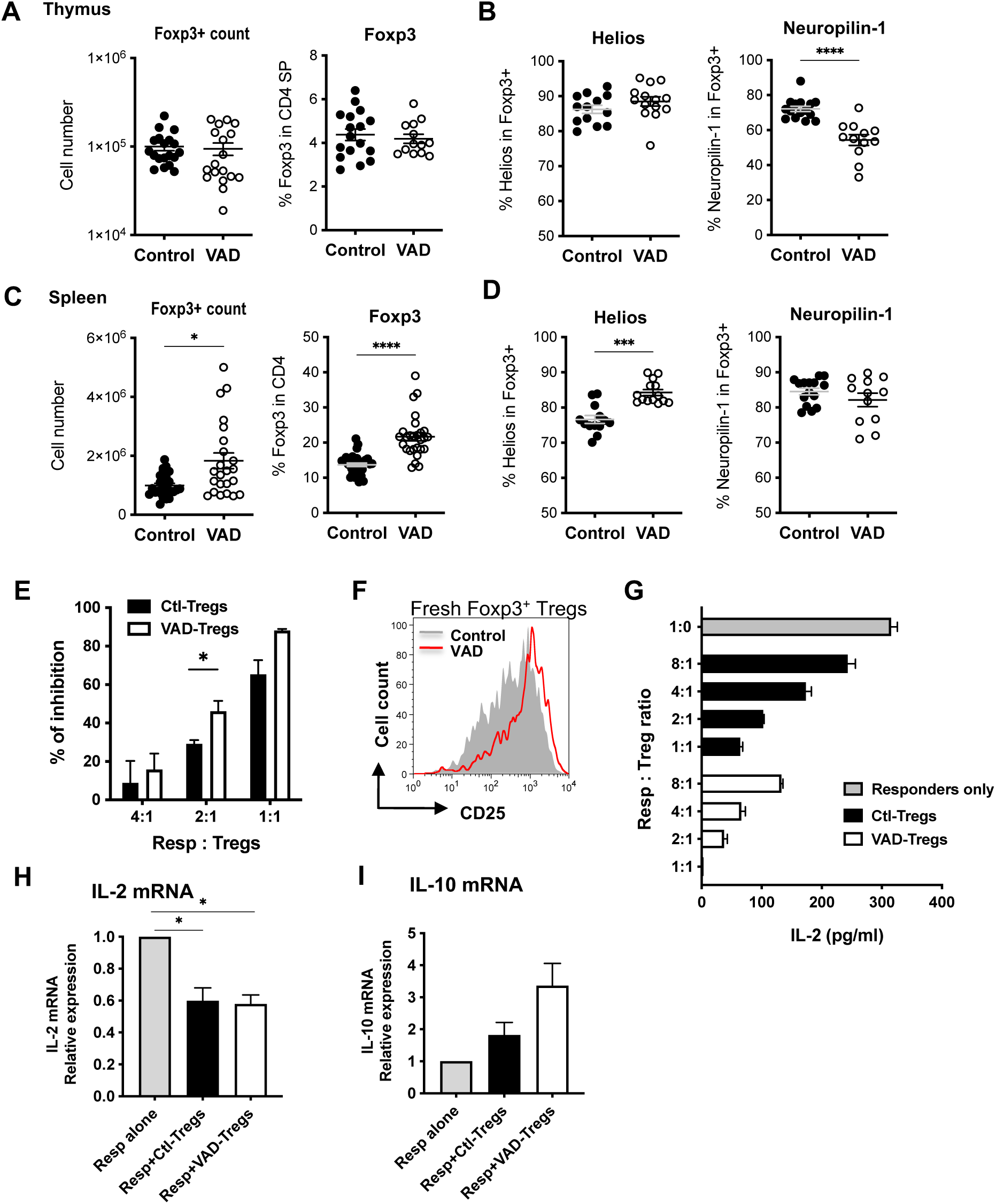
Tregs are present in the thymus and periphery in VAD mice and have regulatory functions. (**A-D**) Absolute cell number and profiles of CD4^+^Foxp3^+^ cells between control and VAD mice in thymus (A) and spleen (C). The frequency of Helios^+^ and Neuropilin-1^+^ cells in Foxp3^+^ cells in the thymus CD4SP cells (B) and spleen CD4^+^ cells (D). (**E-I**) Treg function from control and VAD mice was evaluated by in vitro suppression assays. (**E**) Inhibition of proliferation (anti-CD3 stimulation). Compiled data of 3-4 independent experiments. (**F**) CD25 expression on fresh splenic CD4^+^Foxp3^+^ Tregs. (**G**) IL-2 concentration in culture supernatants was analyzed by ELISA. (**H**, **I**) Responder to Tregs ratio was 2:1. CD4^+^GFP^-^ responder cells were sorted out 36 h after co-culture and analyzed for IL-2 (**H**) and IL-10 (**I**) gene transcription. Compiled data of 3 experiments. \**P* <0.05, \*\*\**P* <0.001, \*\*\*\**P* <0.0001, by t-test.

Functional evaluation of Treg cells sorted from VAD vs. control mice in a standard in vitro co-culture assay with naïve responder T cells revealed that Treg cells from VAD mice were moderately, but statistically significantly, more inhibitory in suppressing proliferation of responders at the 1:2 Treg-to-responder ratio than Tregs cells isolated from control mice (Fig. 2E). Inhibition of IL-2 production and depletion of available IL-2 by Tregs is an important parameter for Treg suppressive function [40]. Freshly isolated VAD Tregs had a higher expression of high affinity IL-2R by CD25 staining, suggesting that they could consume more IL-2 from the cultures than VitA-sufficient Tregs (Fig. 2F). In the Treg-responder co-cultures, Tregs from both control and VAD mice efficiently reduced IL-2 level in a dose-dependent manner, but supernatants from cultures with VAD-Tregs contained less free IL-2 at all Treg-to-responder ratios (Fig. 2G). To address the reason for this at a more mechanistic level, Tregs and responders after 36 h co-culture were separated by sorting for GFP-positive (Treg) and GFP-negative (responder) cells, and were analyzed separately. Results showed that responders that had been co-cultured with VAD or control Tregs had similarly suppressed expression levels of IL-2 mRNA (Fig. 2H). However, responders with VAD Tregs expressed higher IL-10 mRNA (Fig. 2I). Taken together, these data suggest that Tregs from VAD mice are at least as potent functionally as control Tregs in suppressing activation of naïve responder T cells, and that they may do so by consuming more IL-2 and enhancing IL-10 expression in the responders.

### VAD mice fail to develop EAU with active immunization

To evaluate the consequence of VAD on induction of autoimmune uveitis, VAD or control mice were immunized with an IRBP peptide to induce EAU in disease-susceptible strain B10.RIII. Retinol levels in the eye of VAD mice at 9 weeks of age confirmed VitA deficiency at the time of EAU induction (Fig. 3A). Clinical examination of the fundus revealed that VAD mice developed minimal or no disease scores even 21 days after immunization, a time when disease in controls had already peaked a week earlier (Fig. 3B). Histopathology demonstrated severe retinal destruction in the control mice, whereas retinal structure in VAD mice remained largely normal (score 0) and indistinguishable from unimmunized mice (Fig. 3C).

**Figure 3.**
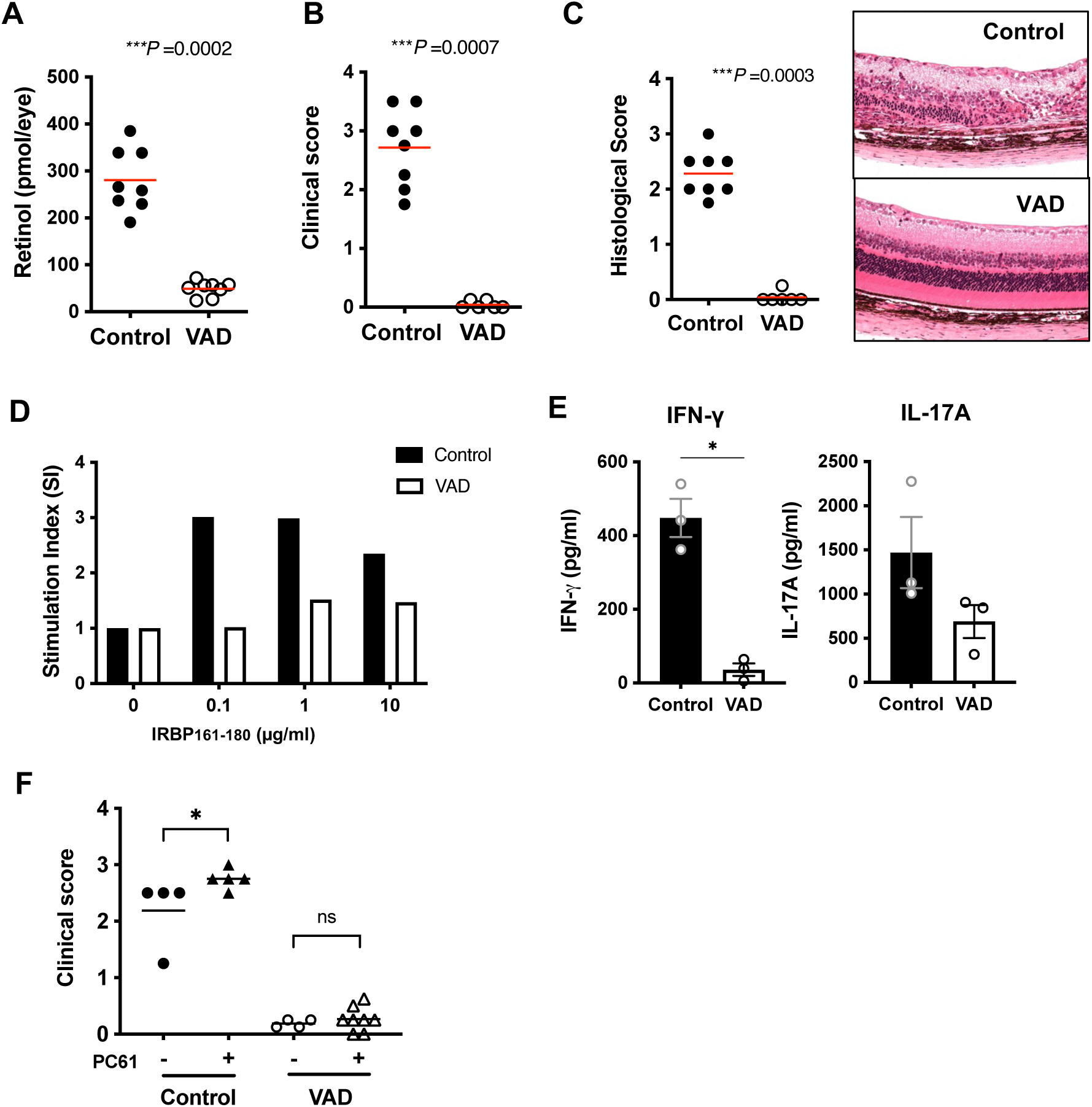
VAD mice fail to develop EAU and show impaired effector T cell functions. B10.RIII mice were immunized with IRBP161-180 in CFA to induce EAU between 9 and 12 weeks of age. (**A**) Retinol levels in the eye of VAD mice. (**B**) Disease was scored by fundoscopic examination between day 17 and 21 for the final time point. One representative experiment of 3 is shown. (**C**) Histology scores (left) and representative H&E slides (right) on day 21. (**D**) Recall antigen-specific splenocyte proliferation in response to IRBP161-180 stimulation. One representative of 5 experiments. (**E**) Effector cytokine production in response to in vitro 48 h recall stimulation with 100 µg/ml IRBP161-180 was measured by ELISA from splenocytes at day 21 post immunization. Data from 3 experiments. (**F**) Depletion of Tregs by anti-CD25 (PC61) did not augment disease in VAD mice. Clinical scores at day 20 were shown. One representative of 4 experiments.

To examine the underlying cellular mechanism, we evaluated lymphocytes from spleens of EAU-induced VAD and control mice for antigen-recall responses in terms of proliferation and cytokine production. Antigen-specific proliferation in lymphocytes from VAD mice was decreased compared to those from control mice (Fig. 3D). Antigen-specific secretion of IFN-γ and IL-17A in the culture supernatant was lower in the VAD group than the control group (Fig. 3E). These data suggest that priming of CD4^+^ T cells from VAD mice had been impaired and therefore these mice are resistant to EAU induction and fail to acquire effector functions.

We next asked whether the impaired priming and acquisition of effector function by VAD CD4^+^ T cells might be the consequence of Treg activity, which appeared to be moderately enhanced in VAD mice (Fig. 2). To address this, mice were depleted of Tregs with anti-CD25 mAb (clone PC61) before immunization and were given a reduced immunization regimen that would be amenable to enhancement by Treg absence. Although PC61 treatment efficiently reduced the CD25^high^Foxp3^+^ Treg frequency in both control and VAD mice (data not shown), disease was enhanced only in VitA sufficient control mice, and VAD mice remained resistant to EAU (Fig. 3F). These results suggest that the defect in acquisition of effector function by CD4^+^CD25^-^ T cells of VAD mice was independent of Tregs.

### Autoimmune tissue damage and inflammation is maintained in a spontaneous model of uveitis when onset of disease precedes onset of VAD

Dietary depletion of VitA as used in the current study results in substantial VitA deficiency by 5 weeks of age and in almost complete deficiency by 9 weeks of age (Fig. 1B) [41, 42]. R161H mice express IRBP-specific TCR and serve as a model of spontaneous autoimmune uveitis. These mice express a transgenic TCR specific for IRBP_161-180_, a major epitope on B10.RIII mice, with a high proportion of their T cells, and spontaneously develop ocular inflammation. First signs of ocular inflammation are apparent in unmanipulated R161H mice around 3–4 weeks of age and disease develops gradually to reach a peak at around 2-3 months [23]. This means that, unlike in the induced EAU model, in which immunization can be done under conditions of full VAD, autoimmunity starts under partial VAD, well before full VAD is reached at 9 weeks of age on the VAD diet [41].

R161H mice were reared on the VAD diet and disease development was monitored by weekly fundus examination. The serum retinol levels confirmed the VitA deficiency in R161H on the VAD diet (Fig. 4A). The uveitis development was comparable in R161H mice on the VAD diet and control diet before 9 weeks of age, when VAD status is still progressing and partial. After 9 weeks of age, the disease scores of R161H-VAD mice were significantly lower than control mice (Fig. 4B). Histological examination revealed a trend towards lower scores in VAD mice (at 12-14 weeks of age), but statistics did not reach a significance (*P*=0.0554). To assess whether antigen presentation by dendritic cells to prime T cells was affected by VAD, highly naïve IRBP-specific T cells were co-cultured with dendritic cells from spleen of WT- or R161H-VAD mice in vitro in the presence of IRBP peptide. Antigen-specific T cell activation judged by early CD69 expression appeared intact in VAD dendritic cell co-culture (Suppl. Fig. S2A, B). Furthermore, in vitro antigen-specific polarization towards Th1 or Th17 was not impaired in R161H-VAD T cells (Fig. 4D). Intriguingly, Treg induction in vitro, as judged by the frequency of Foxp3^+^ cells, was significantly lower in R161H-VAD T cells. However, under the same Treg-inducing condition, more R161H-VAD T cells differentiated into an IL-10 producing population (Fig. 4E). Tregs isolated from R161H-VAD mice were functionally suppressive to naïve responders from either control or VAD mice (Suppl. Fig S3A), similarly to polyclonal Tregs from WT mice (Fig. 2E). However, they were not able to suppress effector/memory T cells (Suppl. Fig. S3B). IL-2 production in the culture supernatant from these suppression assays showed that, in the presence of Treg in the co-culture, IL-2 gene expression and production were decreased in the responder T cells (Suppl. S3C, D). These data indicate that effector function that had been acquired prior to onset of VAD can continue to be expressed and maintained as the mice progress to become full VAD.

**Figure 4.**
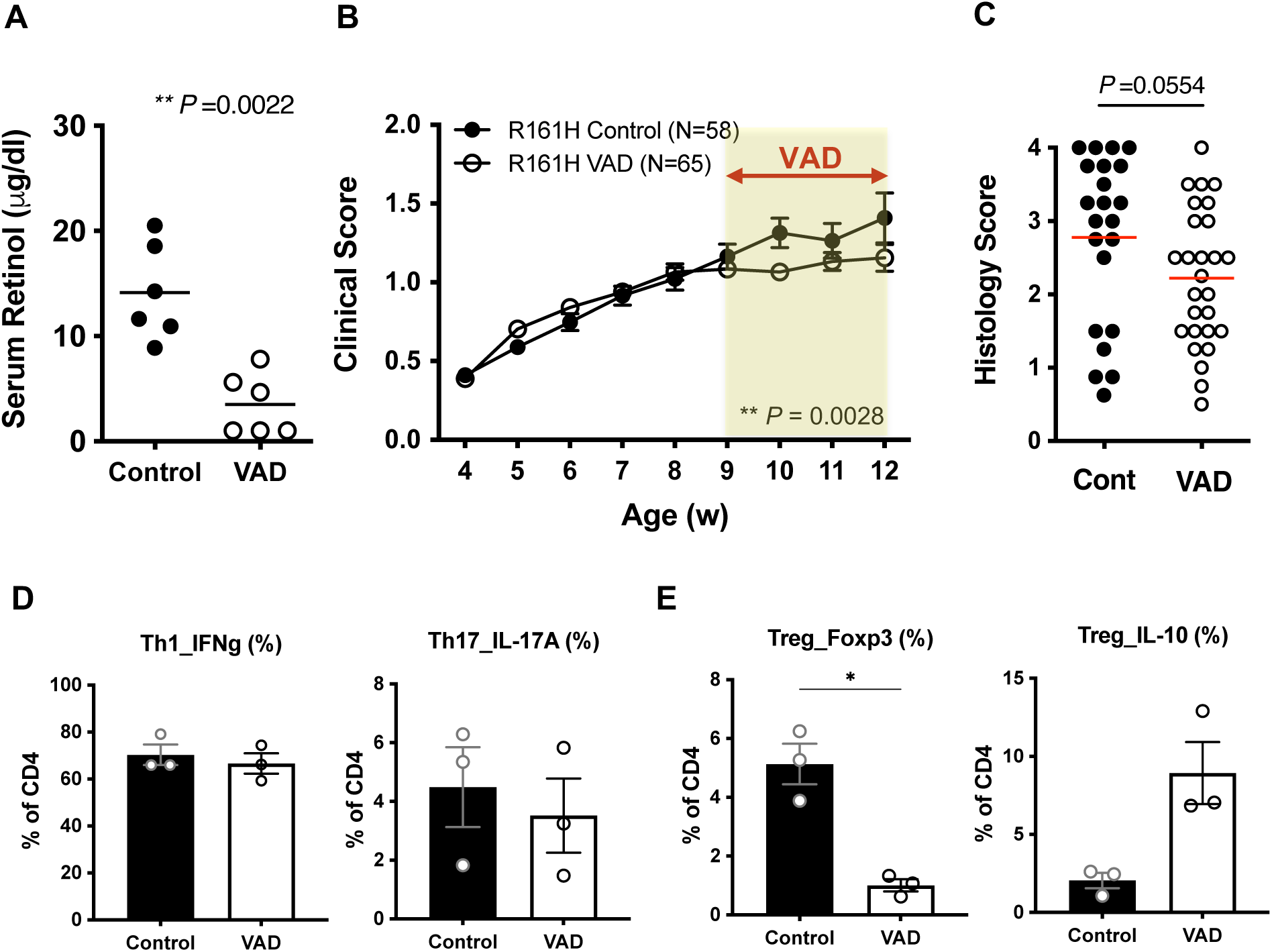
Spontaneous autoimmune uveitis develops in R161H mice in which disease onset precedes onset of VAD. (**A**) Serum retinol levels of R161H mice on control or VAD diet at 11 weeks old. (**B**) Fundoscopy scores of spontaneous uveitis on control or VAD diet. Combined data of 4 experiments. (**C**) Histology data of 12-14 weeks old R161H mice on control or VAD diet. Combined data of 3 experiments. (**D,E**) Purified naïve CD4^+^ T cells from R161H mice on control or VAD diet were polarized under Th1, Th17 (**D**) or Treg (**E**) conditions in the presence of p161-180 (1 µg/ml) and CD11c+ APC from WT mice for 4 days. One representative experiment of three.

### Partial VAD is permissive to in vivo acquisition of effector function and development of autoimmune disease

We next asked whether partial VAD conditions (i.e., VitA insufficiency, VAI), as opposed to full VAD, can also be permissive to immunization-induced EAU. We induced EAU in B10.RIII WT mice on the VAD diet at 5 weeks of age, when the serum retinol levels are still partial (∼50% of controls, Fig. 5A) and compared the phenotype to age-matched control mice on VitA-sufficient diet. In a sharp contrast to the results of full VAD mice, VAI mice developed EAU similarly to control mice in both clinical scores and histopathology (Fig. 5B, C). While antigen-specific recall proliferation was unchanged (Fig. 5D), antigen-specific secretion of IFN-γ and IL-17A appeared lower from the VAI cells than the control cells (Fig. 5E).

**Figure 5.**
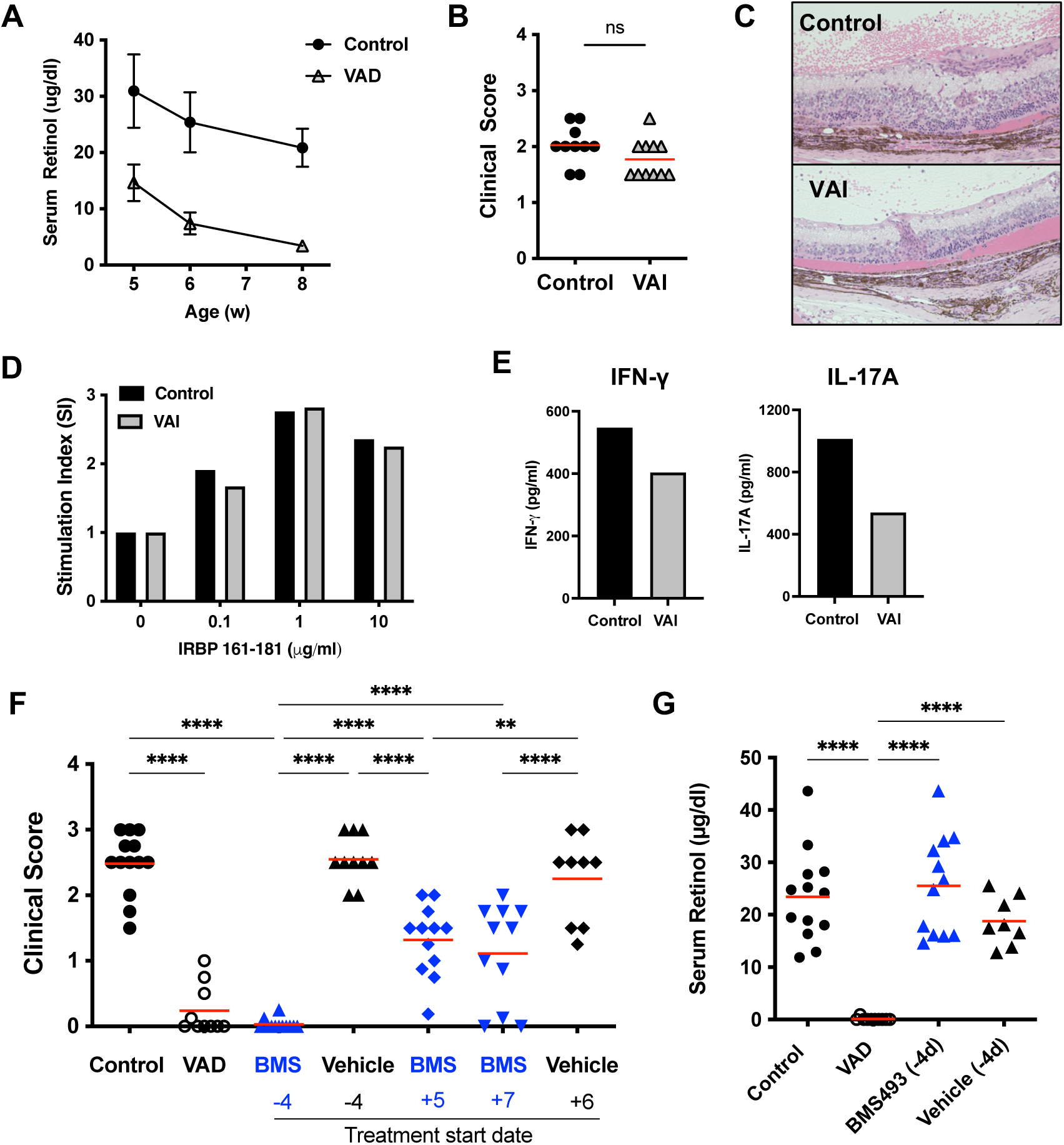
VitA insufficiency (VAI) is permissive to in vivo acquisition of effector function and development of disease. (**A**) Serum retinol levels in VAD and control mice (N=3-6/time point). (**B**) Mice with partial VAD, i.e, VitA insufficiency (VAI) develop as severe disease as control mice when immunized at 5 weeks of age with IRBP161-180 in CFA. Clinical scores evaluated by fundoscopy on day 19 or 22 in 2 experiments. (**C**) Representative histology pictures of a control or VAI mouse. (**D,E**) Antigen-specific recall proliferation and cytokine production at 48 h stimulation with IRBP161-180. (**F, G**) Depleting VitA at the the time of priming (day -4) but not during the effector phase (day 5 to 7) of the immune response prevents the disease development. Ten weeks old mice were immunized with IRBP161-180 in CFA. Daily doses of either vehicle (DMSO) or pan-RA receptor inhibitor, BMS493, were given intraperitoneally. Disease was scored by fundoscopy. Data are combined from 2 independent experiments. Each point represents data from an individual mouse. (**F**) Serum retinol was measured on day 18 (or 19) post immunization. \*\*\*\**P* <0.0001.

As an alternative way to make VAD or VAI mice, we treated mice on the normal chow with daily injections of BMS493, pan-RA receptor inhibitor [27], to block the RA signaling. This method allows us to control the timing of VAD induction either before or after immunization for EAU: BMS493 treatment starting 4 days prior to EAU immunization permits the depletion of VitA at the time of priming, and the treatment starting between day 5 and 7 of EAU permits the depletion of VitA during the effector phase. While BMS493-treated mice from the priming stage (-4) were almost completely resistant to EAU, similar to full VAD mice, those treated from the effector phase (+5, +7) developed EAU and were only partially resistant (Fig. 5F). As the target of the treatment is the RA receptor, serum retinol levels were not affected in mice treated with BMS493 (Fig. 5G). These results support the data from R161H and VAI EAU models that autoimmune disease can develop in the partial VAD environment.

### Induction of EAE is impaired in both VAD and VAI mice

As another and complementary model of CNS autoimmunity, we induced EAE in B10.RIII mice that had been made VAD or VAI, following an optimized immunization regimen with the whole MBP protein [32]. VAD mice with almost complete depletion of serum retinol were essentially resistant to EAE (Suppl. Fig. S4A, B). The resistant phenotype to EAE in VAD mice is in line with, and supports, the EAU results. However, VAI mice immunized for EAE at 5 weeks of age were also significantly resistant to EAE compared to VitA-sufficient mice (Suppl. Fig. S4C, D), whereas VAI mice immunized for EAU at the same age developed disease similarly to controls (Fig. 5B). Nevertheless, VAI mice were more susceptible to EAE than VAD mice which developed no disease at all (Suppl. Fig. S4B, D).

### Supplementation of VitA to VAD mice restores development of EAU

We wanted to explore whether re-feeding VitA to VAD mice can reverse unresponsiveness to EAU induction. After inducing VAD status through dietary deprivation, we replaced the VAD diet with the VitA-sufficient control diet for at least 3 more weeks before EAU immunization. Serum retinol levels confirmed their recovery from the VAD status at the end of the experiment (day 16 post-immunization) (Fig. 6A). Upon induction of EAU, ex-VAD mice developed disease at a severity similar to control mice that had been on VitA-containing control diet for the entire time. VAD mice that continued on the VAD diet throughout the period (>13 weeks) remained almost completely resistant to EAU (Fig. 6B, C). Antigen recall lymphocyte response was improved in ex-VAD mice compared to that in VAD mice (Fig. 6D). These results indicate that although VitA is a critical dietary component that maintains immune responsiveness, immunocompetence is rapidly restored as VitA is replenished.

**Figure 6.**
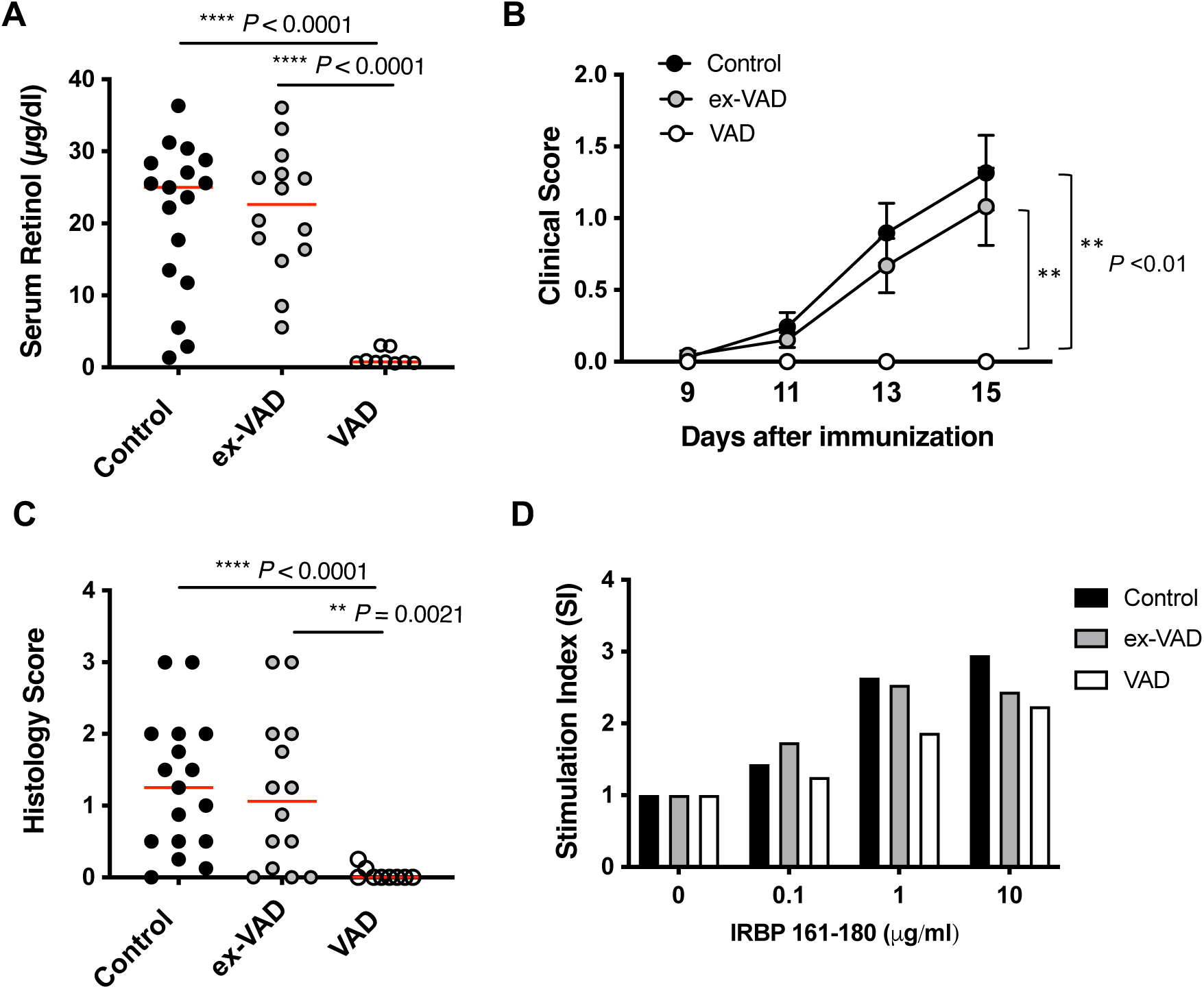
Supplementation of Vit A to VAD mice restores development of EAU. B10.RIII mice were made VAD through dietary means until 12-13 weeks old and recovered on VitA-sufficient diet for at least 3 weeks before immunized with 5 µg IRBP161-180 in CFA. (**A**) Serum retinol levels on day 16 post-immunization. (**B**) Disease was scored by fundoscopic examinations. Average scores of each group at different time points. (**C**) Histology scores on day 16. Combined data from 2 independent experiments (A-C). (**D**) Recall antigen-specific splenocyte proliferation at 48 h. One representative experiment of two is shown. \*\**P* <0.01, *** *P* <0.001 by One-way ANOVA.

### Effector function of already-polarized T cells is poorly controlled in the VAD host

The marginal effect of VAD on spontaneous uveitis in R161H mice suggested that previously acquired effector function can be maintained and expressed in the VAD environment. To address this under more controlled conditions than the “black box” that surrounds spontaneous development of autoimmunity, we employed an adoptive transfer EAU model, using in vitro polarized effector Th17 or Th1 cells specific to retina.

To test the behavior of Th17 cells, R161H T cells from VitA-sufficient donors were polarized towards the Th17 lineage (Fig. 7A) and infused into CD90.1-congenic VAD recipient mice at >9 weeks of age. Onset of EAU induced with adoptively transferred Th17 cells occurs around 7 days after transfer. The VAD recipients infused with R161 Th17 cells showed a trend towards higher disease scores on day 7 than VitA-sufficient control recipients, and by day 10 overall active inflammation in the eye declined (Fig. 7B). Histology examination at the end point (day 10-11) confirmed that VAD recipients developed significantly more severe disease than control recipients (Fig. 7C). As a complementary approach, we performed the adoptive transfer of Th17-polarized cells into BMS493-treated “VAD” recipients vs vehicle-treated recipients. Th17 cells again induced significantly higher disease scores in BMS493-treated recipients than in control recipients (Fig. 7D).

**Figure 7.**
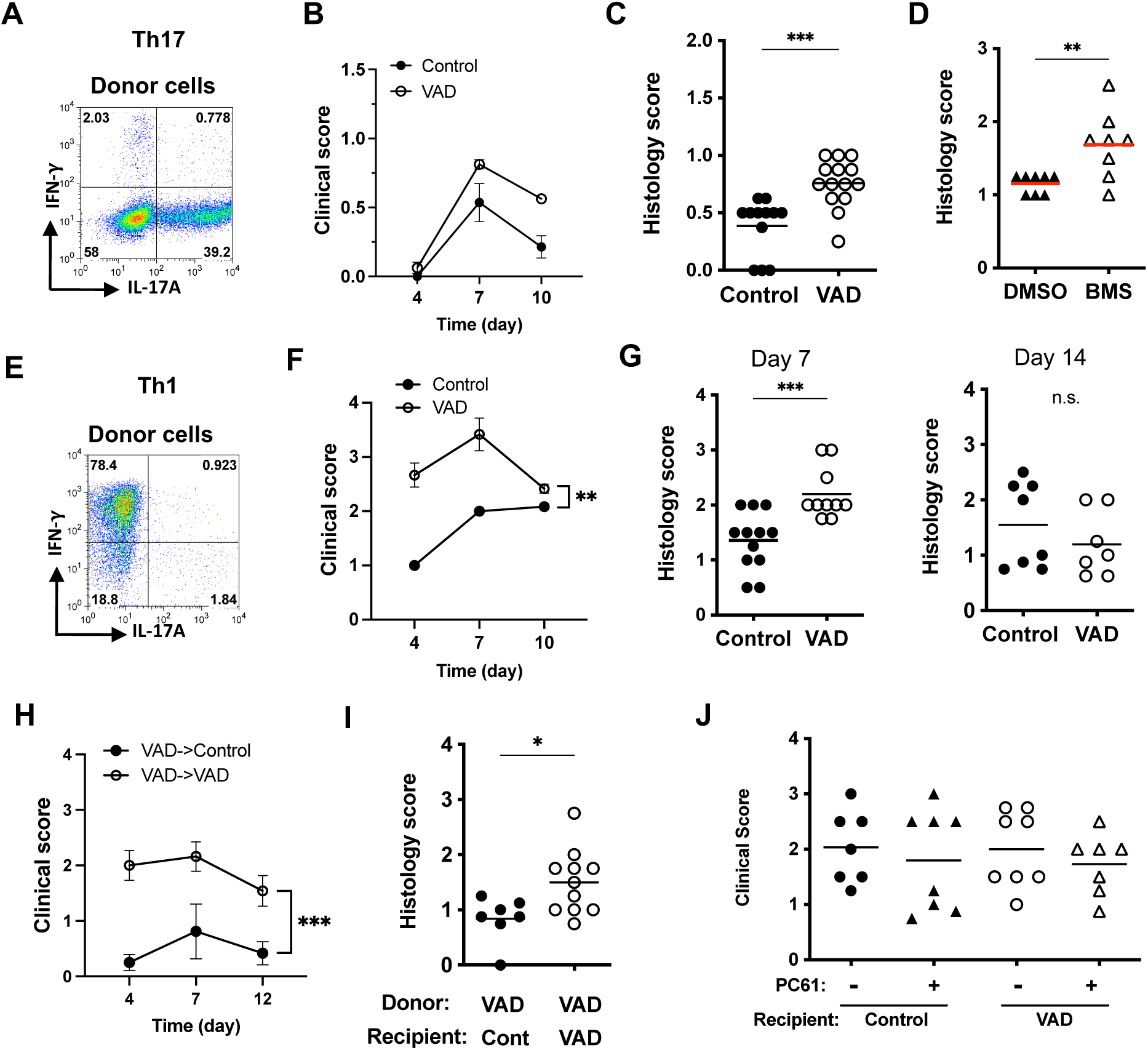
Pathogenic effector cells generated under Vit A-sufficient conditions are poorly controlled in the VAD host. Peripheral lymph node cells from VitA-sufficient R161H mice were polarized towards Th1 or Th17 and used as donor populations and adoptively transferred into control or VAD recipients, or DMSO- or BMS493-treated recipients. (A) Confirmation of Th17 polarization before adoptive transfer. (B) Clinical scores of control or VAD recipient mice post-transfer of Th17 cells. (C) Histology scores at day 10-11 post-transfer of Th17 cells in VAD and control recipients. Data combined from 2 experiments. (D) Histology scores at day 10 post-transfer of Th17 cells in DMSO- or BMS493-treated recipients. (E) Confirmation of Th1 polarization before adoptive transfer. (F) Clinical scores of recipient mice post-transfer of Th1 cells. (G) Histology scores on the 2 different end points, day 7 and day 14, in VAD and control recipients of Th1 cells. Each graph shows combined data from 2 experiments. (H) VAD R161H cells were polarized towards Th1 and adoptively transferred into VAD and control recipients. Clinical scores of control or VAD recipient mice post-transfer. (I) Histology scores at day 10 post-transfer in BMS493-treated recipients. * *P* <0.05, ** *P* <0.01, *** *P* <0.001. (J) Histology scores at day 11 post-transfer in VAD and control recipients treated or not with PC61. Data combined from 2 experiments.

To test the behavior of Th1 cells, we performed adoptive transfer of in vitro polarized Th1 cells (Fig. 7E). Of note, Th1 effector cells from R161H mice are more pathogenic than Th17 effector cells [23]. Onset of EAU transferred with Th1 cells is typically on day 4 after transfer, and disease severity is higher than that induced by Th17 cells. R161H Th1 cells induced significantly more severe disease in VAD recipients than in VitA-sufficient recipients (Fig. 7F). Histology scores on day 7 and 14 confirmed the clinical scores by fundoscopy on day 7 and 10, respectively (Fig. 7G). We also tested how R161H Th1 cells generated from VAD donors express their pathogenicity in VAD vs. control recipients. Similarly to Th1 cells generated from VitA-sufficient mice, VAD Th1 cells induced significantly higher disease in VAD recipients than in control recipients (Fig. 7H). Thus, the poorer control of autoimmune effector T cells by VAD recipients was common to Th17 and Th1 effector responses.

Lastly, to examine the possible contribution of Tregs during the adoptive transfer EAU, we depleted Tregs from the recipient mice by PC61 treatment before adoptive transfer of in vitro-polarized Th1 cells. PC61 treatment showed no effects on disease scores regardless of the VitA sufficiency status of the recipients (Fig. 7J). These results indicate the poor control of activated effector cells by VAD recipients suggests that despite the ability of VAD Treg cells to suppress activation of retina-specific (but naïve) T cells *in vitro*, they were less effective in controlling antigen-experienced effector T cells *in vivo*.

### VitA deficiency compromises ocular immune privilege

Peripheral effects of VitA on immunity would presumably be shared by other autoimmune diseases. The eye, however, is in the unique situation of having a very high local concentration of free retinoids, which are part of the chemistry of vision, but also participate in the phenomenon of ocular immune privilege [19, 20]. Of these, not only RA (all-trans retinoic acid), but also all-trans retinal and possibly all-trans retinol affect Treg induction [43, 44]. The effect of these metabolites, together with TGF-β that is also present in ocular fluids, is to “disarm” retina-specific T cells that may find their way from the circulation into the eye as a result of trauma or vascular abnormality, and prevent them from initiating inflammation [20]. As shown in Fig. 3A, eyes of VAD mice have a drastically reduced level of retinoids. To examine the fate of retina-specific T cells entering the eye of VAD mice, we injected sorted CD4^+^CD44^low^Foxp3^-^ naïve IRBP-specific CD4^+^ T cells from VitA sufficient R161H donor mice into the vitreous cavity of VAD or control recipients. As we reported previously, naïve R161H cells injected into a healthy eye do not induced disease and 20-30% of them were converted to Tregs [20]. However, VAD recipients showed higher clinical scores than control recipients, that showed minimal inflammation, after 6 days of injection (Fig. 8A). This indicates that injected naïve R161H cells persisted more in the eyes of VAD mice and caused stronger inflammation than they did in control recipients. In support of this, IFN-γ- and IL-17A-producing CD4+ donor T cells were markedly higher, whereas Treg conversion was reduced by 50% compared to controls, in eyes of VAD recipients (Fig. 8B). While these naïve T cells came from a VitA sufficient donor, explaining their ability to readily acquire effector function in the VAD environment, these results demonstrate that ocular immune privilege has been compromised in the VAD host.

**Figure 8.**
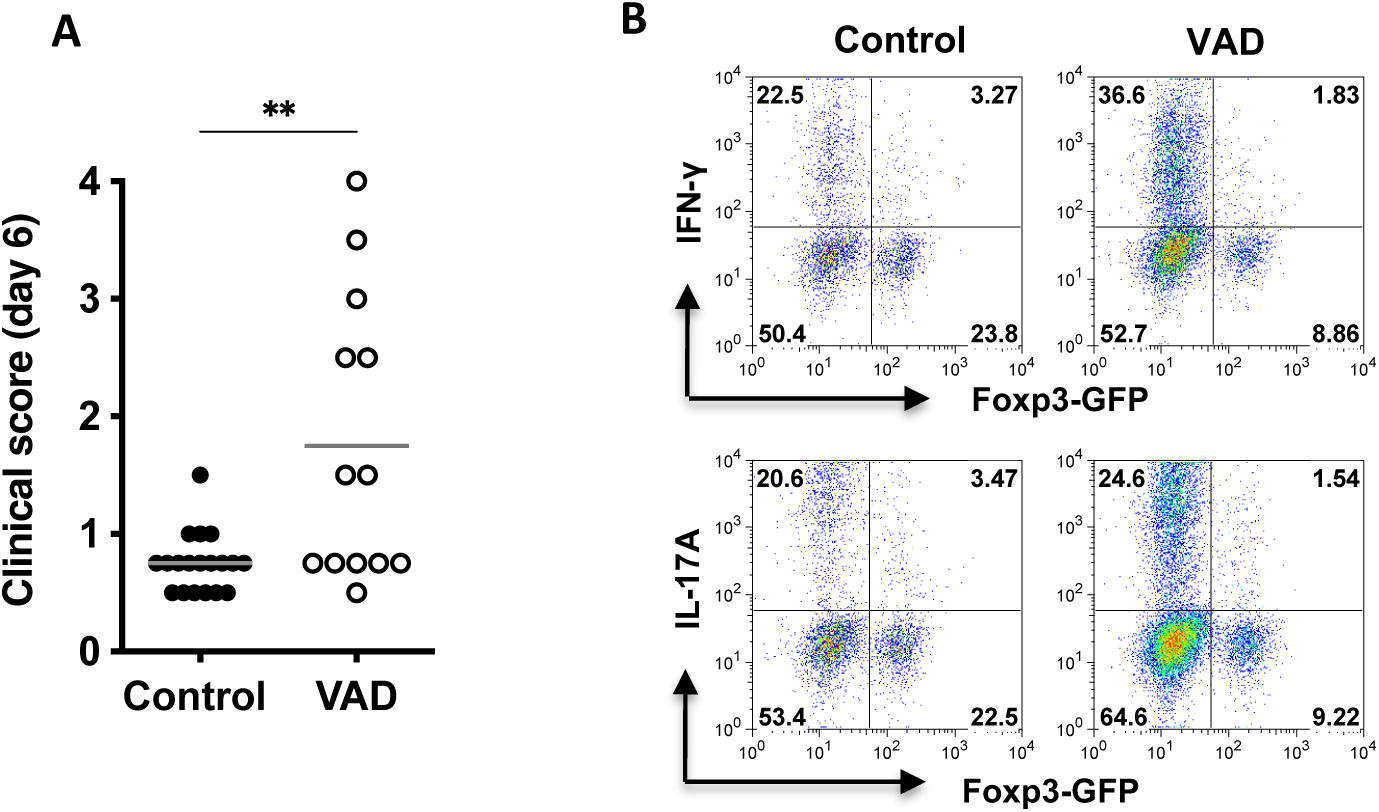
Naïve IRBP-specific T cells introduced into the eye of VAD mice became effector rather than regulatory cells and caused inflammation. CD4^+^CD44^low^Foxp3^-^ naïve CD4^+^ T cells from VitA sufficient R161H-Foxp3^GFP^ donor mice were sorted and intravitreally injected into the healthy eyes of control or VAD hosts. **(A)** Clinical scores for ocular inflammation in recipient mice were examined by fundoscopy at day 6 after naïve cell injection. A score of each eye was plotted, and data from 2 experiments were combined. \*\**P*<0.005 by Unpaired *t*-test. **(B)** Intracellular IFN-γ and IL-17 expression in CD4^+^CD90.2^+^ donor T cells from the pooled recipient eyes on day 6. One representative data is shown from 3 experiments.

## DISCUSSION

Recently, it has been demonstrated that VitA deficiency impedes T cell activation and acquisition of effector function necessary for host defense to microorganisms and response to the dietary antigen, gliadin [21, 22]. Our data in the models of CNS autoimmunity are in agreement with this. In the EAU model that targets the neuroretina, which is an experimental equivalent of the human autoimmune uveitis, immunization of fully VAD mice with the uveitogenic antigen (IRBP peptide) did not result in disease induction (Fig. 3). Although Ag-specific T cells were primed, they exhibited deficient lineage commitment with reduced expression of lineage-specific transcription factors and effector cytokines. Although VAD mice have nTregs that are fully functional or even superior to controls in preventing activation of naïve T cells, this activation defect appeared to be independent of Tregs. Notably, lack of VitA has also been shown to alter the dendritic cell subsets in spleen [42] and dendritic cell function in the mucosal immune system [45]. Our results, however, show that dendritic cells in the spleen of R161H or WT mice were able to activate retina-specific T cells (Suppl. Fig. S2). Although we cannot rule out the involvement of dendritic cells in events leading to disease blockade, dendritic cells appear to function normally in our EAU system. Either way, our data support the notion that an autoimmune priming under conditions of VAD would not result in autoimmune disease.

It might be tempting to speculate that the well-recognized rarity of autoimmune and allergic diseases in the 3^rd^ world countries, where VitA deficiency may be prevalent, could at least in part be due to such mechanisms. However, in “real life” complete and protracted VitA deficiency is probably quite rare (and could be lethal), so immune responses could often be initiated under conditions of only partial VitA deficiency. One question is, what would happen if effector function had been acquired under conditions of relative VitA sufficiency, when the individual subsequently progresses to a VitA deficient state. Our results in spontaneously autoimmune R161H mice where effector cells are generated before onset of full VAD, and in conventional VAD mice adoptively transferred with polarized effector cells, indicate that already acquired effector function is maintained under VitA deficiency and may even be enhanced, triggering exacerbated disease, in some cases (i.e. adoptive transfer in Fig. 7). The mechanism(s) involved in maintenance and enhancement of previously acquired effector function in the VAD environment could at least in part be a consequence of the apparent inability of VAD Tregs to control already committed effector cells. Although the nTregs which are present in VAD mice (and were generated before onset of full VAD) appear quite competent to control activation of naïve T cells, control of committed effectors might require the action of induced Tregs, which are not efficiently generated in VAD mice. Consequently, differences in disease outcome under conditions of partial *vs* full VAD could be qualitative, rather than merely quantitative.

In the specific case of the eye, there is a 2^nd^ layer of complexity to be factored in, and that is the effect of VitA deficiency on ocular immune privilege. As part of ocular immune privilege, the inhibitory ocular environment “disarms” errant retina-specific T cells that enter the eye by converting them to Tregs and/or anergic cells [46]. This happens through the combined action of TGF-β, RA and multiple other molecular triggers, and prevents potentially pathogenic cells from acquiring effector function and precipitating uveitis [20]. Thus, RA in the eye plays a dual role, functioning not only in the chemistry of the visual cycle, but also in ocular immune privilege.

Our data indicate that VitA deficiency impedes generation of locally induced Tregs within the eye, and permits emergence of what could be an effector phenotype (Fig. 8), with the caveat that a broader characterization would be required to confirm this. Although the outcome of this activation process could be affected by these T cells coming from VitA sufficient donors, this does not change the conclusion that ocular immune privilege is compromised in VAD hosts. This may also be applicable to the brain, which is also an immune-privileged tissue, and affect diseases such as multiple sclerosis.

In summary, in our CNS autoimmunity model targeting the neuroretina, VitA deficiency prevented priming and acquisition of autopathogenic effector function, apparently independently of its effects on Treg function. Importantly, however, previously acquired effector function was not only maintained, but even enhanced in the VAD host, resulting in exacerbated disease. The latter finding may be due to reduced ability of the Treg compartment of VAD mice, which – although able to prevent T cell priming – is insufficient to control already committed effector T cells. This extends the findings reported in other VAD models. Because complete VitA deficiency in human populations may be the exception rather than the rule, our findings may have implications on induction and expression of autoimmunity in geographical areas where availability of dietary VitA is limiting or fluctuates seasonally.

## Abbreviations

VitA: vitamin A
RA: retinoic acid
VAD: vitamin A deficient/deficiency
Treg: regulatory T cells
IRBP: interphotoreceptor retinoid-binding protein
EAU: experimental autoimmune uveitis
EAE: experimental autoimmune encephalomyelitis

## ACKNOWLEDGEMENTS

The authors thank Dr. Alexander Y. Rudensky (Sloan-Kettering, NY) for the Foxp3^GFP^ reporter mice, and former and current Caspi lab members for assistance and helpful discussions We thank the staff of the NEI Flow Cytometry Core Facility and the NEI Histology Core Facility for cell sorting and for histology slides processing, respectively. This work was supported by NIH/NEI Intramural funding, Project No. EY000184.

