## Supplemental Figures for "Vitamin A is necessary for acquisition, but not for expression or progression, of CNS autoimmunity"

### Suppl. Figure S1

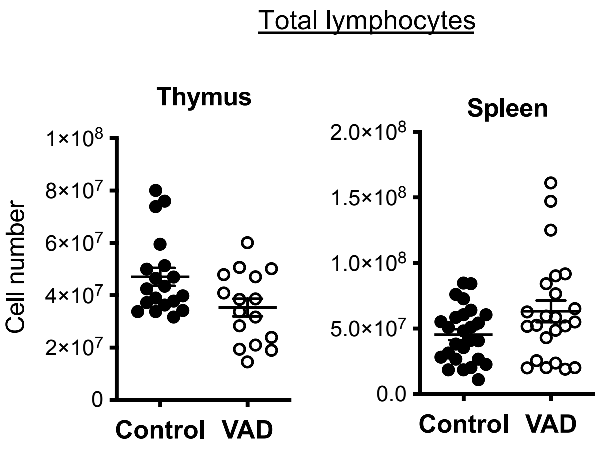


**Suppl Figure 1. Total lymphocytes counts are unchanged in VAD mice.**
Data are compiled from 6 to 8 experiments.

### Suppl. Figure S2

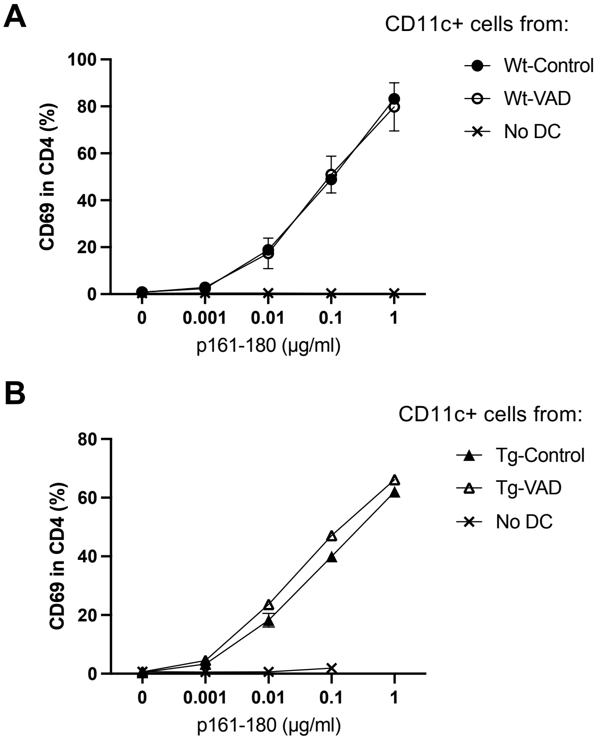


**Suppl. Figure S2. CD11c+ dendritic cells from VAD mice are functionally intact in priming retina-specific T cells.** Dendritic cells were isolated for the CD11c+ populations from spleen of VAD or control mice at 11 wk old or older age. Responder naïve CD4 T cells were purified from R161H Rag2$^{-/-}$ IRBP$^{-/-}$ mice. T cells and DC were co-cultured in the presence of varying concentrations of p161-180 for 20 h. (**A**) CD69 expression on the responder T cells cultured with WT CD11c+ cells. (**B**) CD69 expression on the responder T cells cultured with R161H CD11c+ cells. T cell only (No DC) wells served as negative controls. One representative experiment of 5 with 2-3 mice per group is shown.

### Suppl. Figure S3

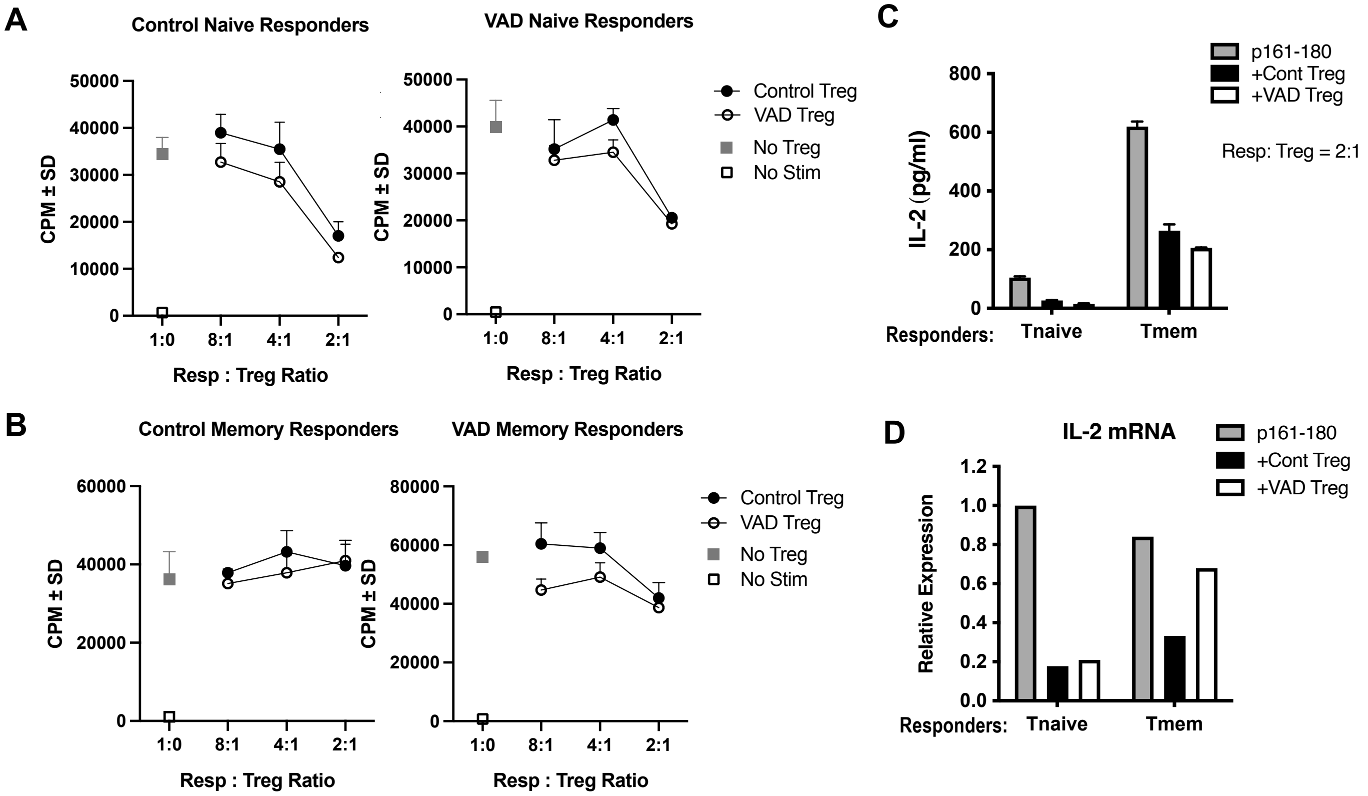


**Suppl. Fig.S3. IRBP-specific Tregs from VAD and control mice have similar suppressive functions.** Treg function from control and VAD R161H mice was evaluated in suppression assays with various numbers of Tregs in the co-culture. Naïve (CD62L$^+$CD44$^{lo}$) or memory (CD62L$^-$CD44$^{hi}$) responder (GFP-) or Treg (GFP+) cells were sorted from R161H-Foxp3$^{GFP}$ mice. Proliferative responses of (**A**) naïve responders from control or VAD mice or (**B**) memory responders from control or VAD mice, were measured in the absence or presence of Tregs with various ratios, cultured with antigen-presenting cells and IRBP161-180 peptide. (**C**) IL-2 concentration in culture supernatants after 18 h co-culture as analyzed by ELISA. (**D**) CD4+GFP- responder cells were sorted out 18 h after co-culture and analyzed for IL-2 mRNA. Relative expression to control Tn (without Treg) was calculated after normalization to respective GAPDH expression. (**C-D**) Responder to Tregs ratio was 2:1.

### Suppl. Figure S4

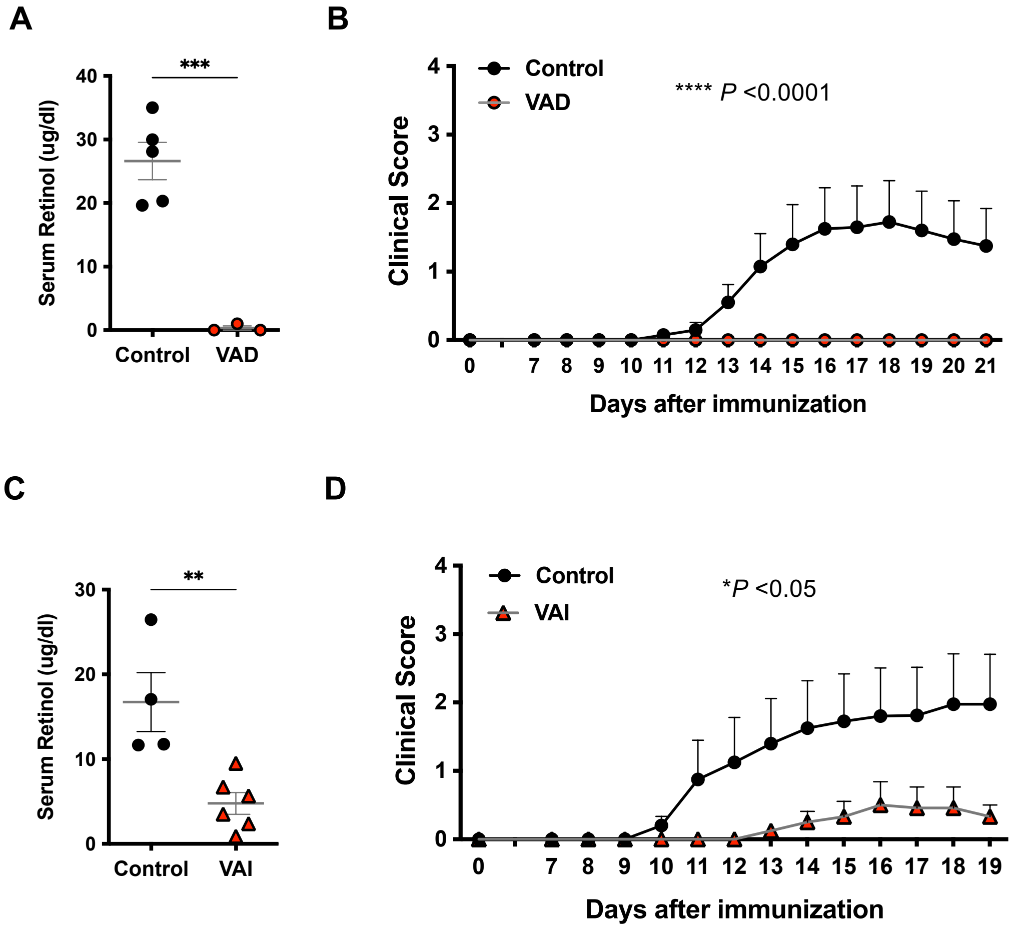


**Suppl Figure S4. Induction of EAE is impaired in both complete and partial VAD mice.** (**A,B**) B10.RIII mice were made VAD through dietary means and immunized at 9 wk of age with 300 µg MBP (whole protein) in 4 mg/ml CFA and two i.v injections of 0.2 µg PTX on day 0 and day 2. N=10 for Control and N=8 for VAD. Compiled data from two experiments. (**C,D**) Partial VAD (VAI) B10.RIII mice at 5 wk of age were immunized as described above. N=5 for Control, N=6 for VAD. One representative of two experiments. (**A, C**) Serum retinol were measured at the end of the experiments. ***P*<0.01, *** *P*<001 by unpaired t test.
